# Retinoic acid signaling suppresses chondrocyte identity during cartilage development and regeneration

**DOI:** 10.1101/2024.05.10.593640

**Authors:** Claire Arata, Sandeep Paul, Simone Schindler, Mathi Thiruppathy, Mackenzie Flath, Dion Giovannone, Zack Hammer, Dev Subramanie, J. Gage Crump

**Affiliations:** Eli and Edythe Broad California Institute for Regenerative Medicine Center for Regenerative Medicine and Stem Cell Research, Department of Stem Cell Biology and Regenerative Medicine, University of Southern California Keck School of Medicine, Los Angeles, CA 90033, USA

**Keywords:** zebrafish, cartilage regeneration, hereditary multiple osteochondromas, growth plate skeleton

## Abstract

Cartilage is a dynamic tissue during development, repair, and disease. Within developing endochondral bones, chondrocytes at growth plate edges transition to osteoblasts, adipocytes, and other connective tissue in the marrow cavity, and in diseases such as Multiple Osteochondromas (MO) chondrocytes form abnormally around growth plates and contribute to ectopic bone. On the other hand, the inability to form new chondrocytes to maintain damaged joints contributes to the high incidence of osteoarthritis. In order to assess the ability of zebrafish to regenerate cartilage, we developed a nitroreductase-based cartilage ablation model. Following ablation at larval to adult stages, we observed new chondrocytes forming around the dead cartilage matrix, with cartilage outgrowths in ablated endochondral bones resembling the exostoses of MO. By generating a perichondrium-restricted *hyal4:GFP* transgenic line, we show that new chondrocytes arise from the perichondrium surrounding the ablated cartilage. In addition, we observe enriched expression of the retinoic acid (RA) synthesis gene *aldh1a2* in the perichondrium following ablation, and the RA degrading enzyme gene *cyp26b1* in newly forming chondrocytes. Consistent with RA signaling suppressing chondrogenesis, treatment with the RA receptor gamma agonist palovarotene, which has been used to treat MO, prevented ablation-induced chondrogenesis. Although we find that BMP signaling is also required for ablation-induced chondrogenesis, we show a distinct role for RA signaling in suppressing *sox9a* expression and chondrogenesis. Moreover, palovarotene resulted in a near complete loss of growth plates in uninjured zebrafish, which was due to the precocious dedifferentiation and exit of chondrocytes from the growth plate. Our findings suggest a common chondrocyte suppressive role of RA signaling within the developing growth plates and regenerating perichondrium, which may explain the growth plate defects observed when using RA agonists to treat MO.

## Introduction

Cartilage is an avascular tissue composed of chondrocytes that produce various collagens and proteoglycans of a specialized extracellular matrix. During endochondral bone formation, chondrocytes are arranged in distinct zones within growth plates, where they transition from resting to proliferative, prehypertrophic, and hypertrophic states as they mature. In both zebrafish and mice, invasion of vasculature into the cartilage coincides with the terminal maturation of hypertrophic chondrocytes, wherein they die or change fate to become osteoblasts, adipocytes, and marrow stromal cells (Giovannone et al., 2019; Long et al., 2022; Mizuhashi et al., 2018; Park et al., 2015; G. Yang et al., 2014; L. Yang et al., 2014). Maturation of the growth plate is tightly regulated by a negative feedback loop involving PTHrP in the resting zone and Ihh in the hypertrophic zone (Karp et al., 2000; Kobayashi et al., 2002; Mak et al., 2008; Minina et al., 2001). However, the transformation of hypertrophic chondrocytes into osteoblasts and other marrow cell types is less well understood.

Compared to bone, cartilage has poor regenerative capacity in mammals. During fetal stages and in children, fractures to the growth plates can heal, albeit with risk of angular deformity depending on the plane of fracture (reviewed in Rodríguez-Merchán, 2005). In certain instances, injury to the perichondrium as well the growth plate can negatively impact the endogenous repair capacity (Sabharwal and Sabharwal, 2018). Later in life, the ability to regenerate damaged cartilage is more limited. One exception is the outer ear, where injury can induce formation of new chondrocytes and fibrocartilage in a condition called auricular hematoma or “cauliflower ear” (Ohlsén et al., 1975; Schonauer et al., 2002). African spiny mice (*Acomys kempi*) but not lab mice (*Mus musculus*) are also capable of regenerating cartilage after ear punches (Seifert et al., 2012), although this involves a distinct type of elastic cartilage from endochondral bones. In contrast, loss of the cartilage lining joints contributes to irreversible damage leading to osteoarthritis, one of the most common debilitations in humans. Articular cartilage in mammals shows only very limited regeneration capacity. In mouse models, variable chondrocyte formation has been reported in a minority of animals after meniscus surgery to induce osteoarthritis (Kania et al., 2020). DBA/1 mice, which have much improved articular cartilage regeneration capacity compared to C57BL/6 mice, still have limited repair in adults (Eltawil et al., 2009). In rabbits, articular cartilage regeneration was observed in young and adolescent animals but did not extend to adults (Wei et al., 1997). In contrast, adult zebrafish can mount a significant regenerative response of jaw joint articular cartilage following joint destabilizing ligament surgery (Smeeton et al., 2022). In skates, a non-bony fish that retains a fully cartilaginous skeleton, cartilage also repairs well after injury in adults, with injury to the perichondrium alone sufficient to induce substantial ectopic cartilage formation (Marconi et al., 2020).

Although mammalian cartilage has limited regeneration potential, there are a number of human diseases characterized by inappropriate formation of cartilage and endochondral bone. These include heterotopic ossification, fibrodysplasia ossificans progressiva, and multiple osteochondromas (MO, also known as hereditary multiple exostoses). MO is caused by heterozygous loss of *EXT1* or *EXT2*, which encode glycosyltransferases involved in synthesis of heparan sulfate proteoglycans (HSPGs) abundant in cartilage matrix (Alvarez et al., 2006; Francannet et al., 2001; Xu et al., 1999). In MO, ectopic cartilaginous growths, or exostoses, form just outside the growth plates of long bones, with these benign tumor-like growths organizing into similar zones to the growth plate and eventually ossifying (Matsumoto et al., 2010). In mice, conditional loss of *Ext1* in populations labeled by *Col2-CreERT*, *Fsp1-Cre*, *Agr-CreERT2*, or *Gdf5-Cre* result in exostoses, suggesting that exostoses may arise from the perichondrium, periosteum, or specialized cells in the Groove of Ranvier (Huegel et al., 2013; Inubushi et al., 2018, 2017; Matsumoto et al., 2010; Mundy et al., n.d.). Exostoses are also seen in zebrafish mosaic for wild-type cells and cells mutant for the *EXT2* homolog *dackel* (Clément et al., 2008). In both mice and zebrafish, loss of HSPGs also leads to recruitment of wild-type cells into exostoses, indicating roles of tissue-wide HSPGs in preventing ectopic cartilage formation.

Both RA and BMP signaling have been linked to the cartilage exostoses in MO. Forced activation of RA signaling by the RA receptor gamma agonist palovarotene (PVO) can prevent exostoses in mouse models of MO, and PVO has received limited approval to treat MO and other disorders of ectopic endochondral bone (Inubushi et al., 2018; Lees-Shepard et al., 2018; Shimono et al., 2011). Similarly, BMP inhibitors prevent ectopic cartilage formation in *Col2-CreERT*; *Ext1^fl/fl^*, *Agr-CreERT2^fl/fl^*; *Ext1^fl/fl^*, and *Fsp1:Cre*; *Ext1^fl/fl^* mouse models of MO (Inubushi et al., 2017; Sinha et al., 2017), consistent with the observation of increased BMP signaling in MO (Huegel et al., 2013; Kawashima et al., 2020; Matsumoto et al., 2010). In fibrodysplasia ossificans progressiva, a rare disease of heterotopic ossification in soft tissues caused by constitutively active BMP signaling, PVO has also shown some success in mouse models (Chakkalakal et al., 2016). While some reports show that PVO treatment can secondarily result in reduced BMP activity in the perichondrium and periosteum (Inubushi et al., 2018), how RA and BMP pathways interact in MO remains unclear.

Despite the success of PVO in suppressing exostoses, severe side effects in adolescents remain a concern. For example, a phase 3 clinical trial of PVO in adolescents was recently halted due to accelerated closure of growth plates (Pignolo et al., 2023). The role of RA signaling in growth plate biology has remained elusive (Koyama et al., 1999; Yoshida, 1999). RA induces transcriptional responses by binding to three types of retinoic acid receptors (RARα, RARβ, RARγ) that form heterodimer complexes with RXRs (reviewed in Moras and Gronemeyer, 1998). In the absence of RA ligand, RARs are also thought to mediate transcriptional repression (Koide et al., 2001; Kurokawa et al., 1995). In mice, RARγ is expressed in the resting and proliferative zones of the embryonic growth plate, RARα is expressed in the perichondrium, and RARβ is expressed broadly at low levels (Williams et al., 2009). Conditional deletion of both RARγ and RARβ or RARα using *Col2a1-Cre* led to reductions in resting and proliferative but not hypertrophic zones, as well as reduced expression of *Aggrecan* and *Sox9* (Williams et al., 2009). However, examination of RARE (Retinoic Acid Responsive Element) reporter mice showed that RA signaling is active primarily in the hypertrophic zone and perichondrium (Schroeder and Heersche, 1998), and treatment of cultured chondrocytes with exogenous RA leads to a reduction in cartilage matrix synthesis (Garcia et al., 2020; Horton et al., 1987; Horton and Hassel, 1985; Williams et al., 2009). These data suggest that RARγ and RARβ may have ligand-independent functions in the resting and proliferative zones to promote chondrogenesis, and then shift to chondrocyte repression when exposed to RA ligand in the hypertrophic zone.

In order to understand whether zebrafish are able to broadly regenerate cartilage throughout their lifetime, here we utilize an animal-wide cartilage ablation model and find that zebrafish can robustly regenerate diverse cartilages through adulthood. The similarity of ablation-induced chondrogenesis to the exostoses seen in MO, particularly around the growth plates of endochondral bones, led us to examine whether common mechanisms underlie the chondrogenic response to cartilage ablation and the reduction of HSPGs in the cartilage matrix. Using a newly generated perichondrium-specific *hyal4:GFP* transgenic line, we show that ablation-induced chondrocytes arise from the perichondrium. We also find that, as with MO, cartilage outgrowths induced by ablation can be inhibited by increasing RA signaling with PVO or all-trans retinoic acid, or by blocking BMP signaling with LDN-193189. Expression of the RA-synthesis gene *aldh1a2* in the perichondrium and the RA-degradation gene *cyp26b1* in new chondrocytes supports a model in which RA-dependent signaling in the perichondrium initiates ablation-induced chondrogenesis and then shifts to ligand-less repression to allow cartilage differentiation. Consistently, inhibition of RA signaling with DEAB also blocks ablation-induced chondrogenesis. Finally, we show that, as in mice and adolescent patients, treatment of uninjured zebrafish with PVO results in abnormal closure of growth plates, with lineage tracing and marker expression revealing that this is due to accelerated exit of chondrocytes from the growth plate into the marrow cavity. Our findings reveal common roles of RA signaling in suppressing chondrocyte identity in the late hypertrophic zone of the growth plate and the ectopic outgrowths seen upon cartilage ablation, thus potentially linking the growth plate closure and exostoses inhibitory effects of PVO in MO patients. In addition, the finding of common mechanisms of MO and ablation-induced chondrogenesis suggest that MO may be considered a disease of misregulated cartilage regeneration.

## Results

### Imperfect regeneration following cartilage ablation in zebrafish

To conditionally ablate cartilage in zebrafish, we utilized a *col2a1a^BAC^:mCherry-NTR* line in which an mCherry and nitroreductase (NTR) fusion protein is expressed under the control of the *col2a1a* locus (Fig. 1a). NTR metabolizes metronidazole (MTZ) into a toxin, allowing cell-autonomous ablation of cells (Askary et al., 2015; Curado et al., 2008). Following a single 24 h treatment of 14 day post-fertilization (dpf) *col2a1a^BAC^:mCherry-NTR*; *col2a1a1^BAC^:GFP* fish with MTZ, we observed reduction of mCherry and GFP fluorescence throughout the face at 16 hours post-ablation (hpa), except for chondrocytes in joints and a subset of the anterior growth plates of the ceratohyal (Ch) cartilages (Fig. 1b). Next, we performed three consecutive rounds of MTZ (24 h on, 24 h off) to ensure complete ablation of cartilage at 14 dpf and performed pentachrome histological staining at 7 days post-ablation (dpa). Compared to DMSO-treated controls, sections of MTZ-treated fish revealed poorly stained cartilages devoid of nuclei, consistent with TUNEL staining at 3 and 5 dpa revealing extensive cartilage cell death (Fig. 1c-e). At 7 dpa, we observed extensive new cartilage around the entire dead shell of the lower jaw Meckel’s (M) permanent cartilage. In contrast, new cartilage outgrowths were localized to regions adjacent to the ablated growth plates of endochondral bone cartilages such as the Ch and palatoquadrate (Pq). In the outgrowths adjacent to Ch growth plates at 3 dpa, BrdU incorporation revealed cell proliferation and *sox9a* expression revealed new cartilage differentiation (Fig. 1f,g). We also observed efficient ablation after a single 24 h treatment with MTZ at juvenile (6-8 weeks post-fertilization (wpf)) and adult (1 year) stages. Alcian blue staining of cartilage in juveniles at 6 dpa revealed new cartilage forming around the entirety of permanent M cartilage and adjacent to the ablated growth plates of Ch and Pq endochondral bones (Fig. 2a-c). At adult stages at 7 dpa, H&E staining and in situ hybridization for *col2a1a* mRNA revealed new cartilage forming around ablated M cartilage (Fig 2d,e). These findings demonstrate that zebrafish are capable of imperfect regeneration of cartilage through adult stages.

**Figure 1:**
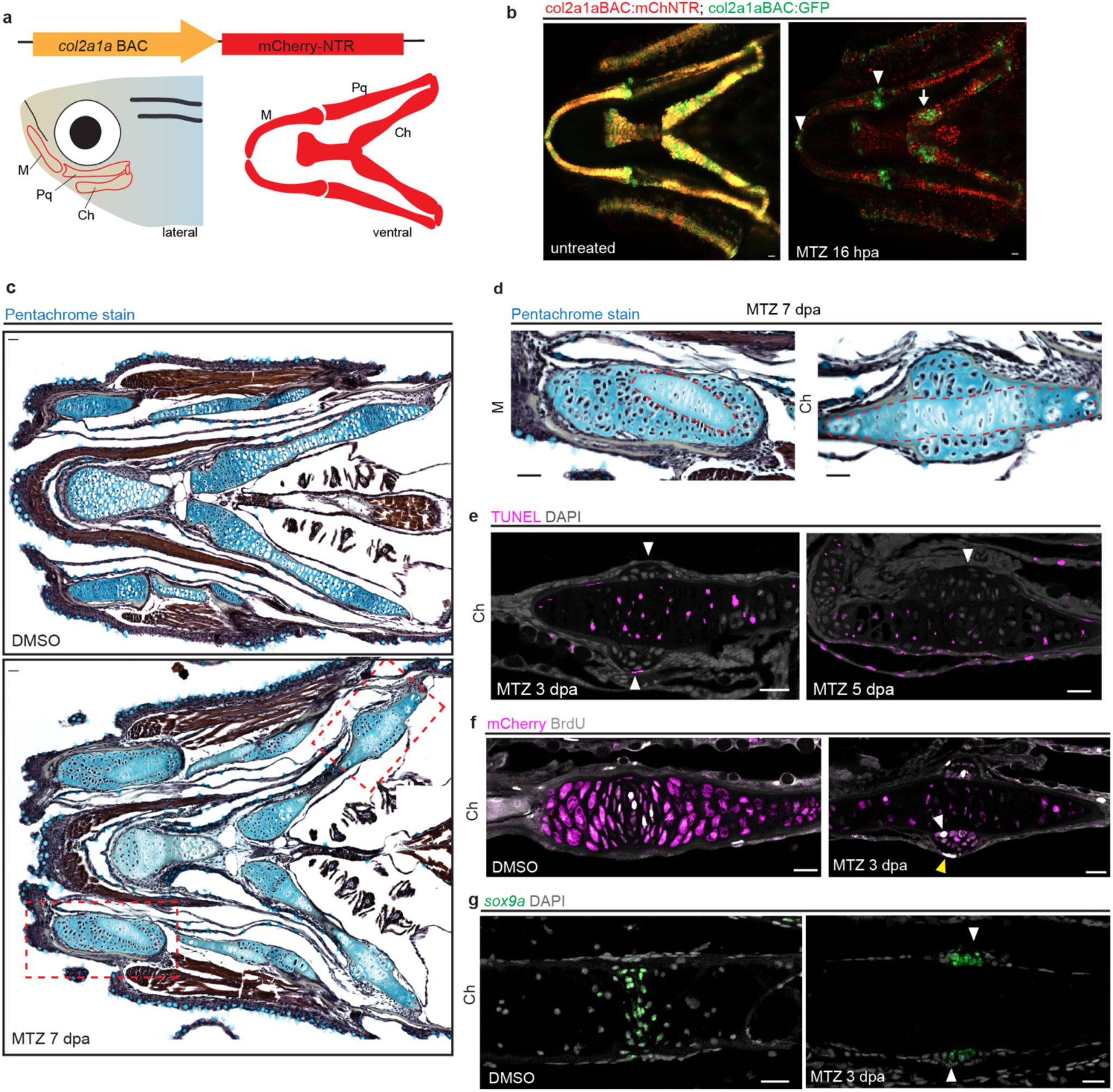
Cartilage ablation and imperfect regeneration in juvenile zebrafish. **a,** Diagrams show Meckel’s (M), palatoquadrate (Pq), and ceratohyal (Ch) cartilages of the zebrafish face that express *col2a1aBAC*:*mCherry-NTR*. **b,** Facial cartilages are labeled by both *col2a1aBAC*:*mCherry-NTR* (red) and *col2a1aBAC*:*GFP* (green) in untreated zebrafish at 14 dpf. Following a single 24 hour MTZ treatment, GFP signal is lost at 16 hpa except for joints (arrowheads) and the anterior Ch growth plates (arrow). **c, d,** Coronal sections of the jaws stained with pentachrome. *col2a1aBAC*:*mCherry-NTR* zebrafish (28 dpf) were treated with DMSO control or three daily doses of MTZ to ablate cartilage, followed by fixation 7 days later. Red boxed regions enlarged in **d** show extensive new cartilage surrounding ablated regions (red dotted lines). **e,** TUNEL staining (magenta) of sections from MTZ-treated *col2a1aBAC*:*mCherry-NTR* fish (21 dpf) shows cell death throughout the Ch cartilage but not the regenerative cartilage (white arrowheads) at 3 and 5 dpa. **f,** BrdU staining (white) of DMSO-treated and MTZ-treated *col2a1aBAC*:*mCherry-NTR* fish (21 dpf) at 3 dpa shows proliferation in chondrocytes (white arrowhead, labeled by mCherry in magenta) and in perichondrium (yellow arrowhead) within new cartilage growths around ablated Ch. **g,** Whereas *sox9a* expression (green) labels Ch growth plate chondrocytes of *col2a1aBAC*:*mCherry-NTR* fish at 21 dpf, only new chondrocytes in outgrowths (white arrowheads) following MTZ treatment express *sox9a* at 3 dpa. Scale bars = 20 um.

**Figure 2:**
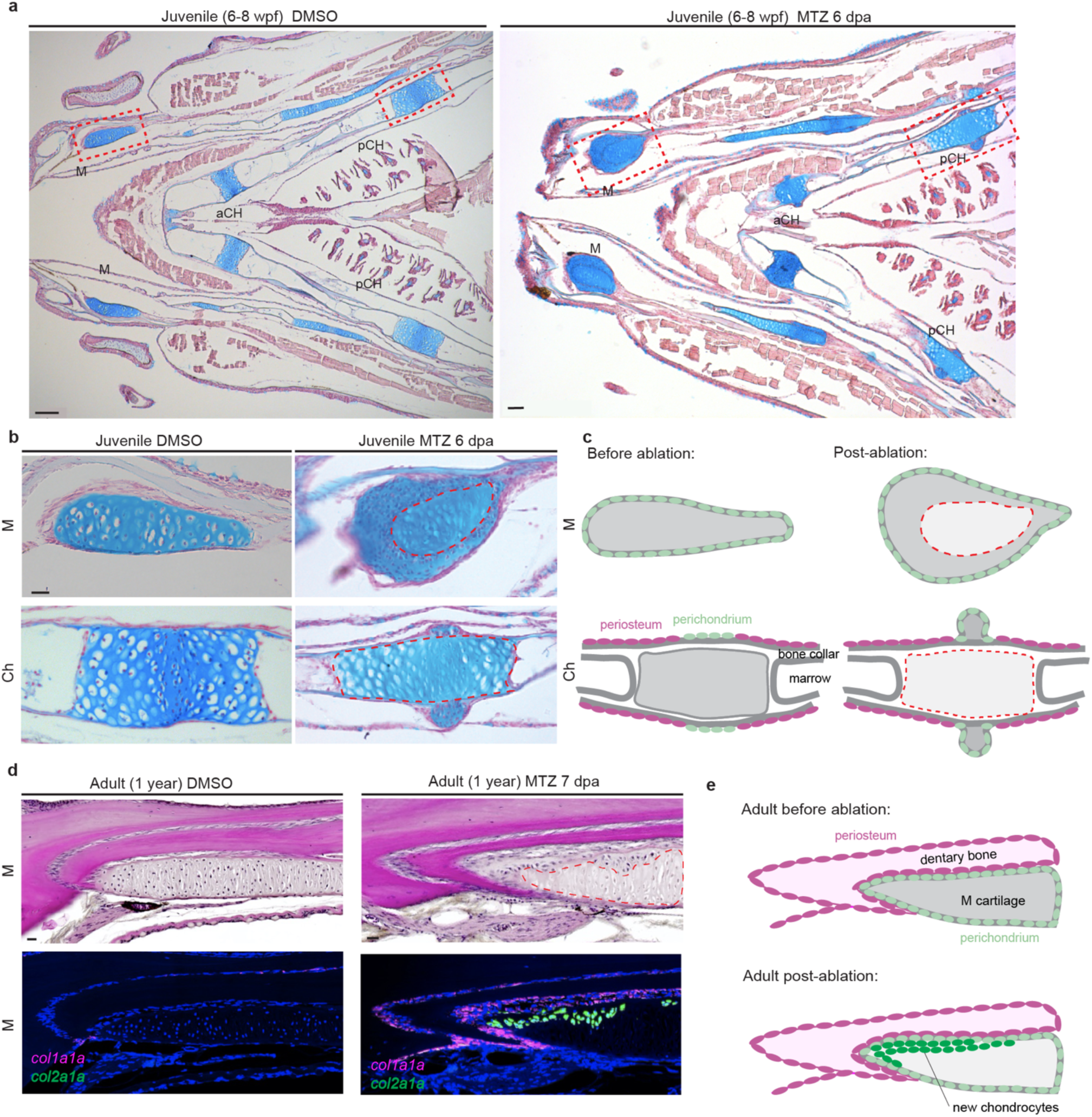
Ablation induces imperfect cartilage regeneration through adult stages. **a, b,** Coronal sections of the jaws stained with Alcian blue and nuclear fast red. *col2a1aBAC*:*mCherry-NTR* zebrafish (6-8 wpf) were treated with DMSO control or MTZ to ablate cartilage, followed by fixation 6 days later. Red boxed regions enlarged in **b** show extensive new cartilage surrounding ablated regions (red dotted lines). **c,** Diagram of the juvenile M and Ch cartilages before and after ablation highlighting new cartilage formation from the perichondrium. **d,** H&E-stained sections of the M cartilages of DMSO or MTZ-treated *col2a1aBAC*:*mCherry-NTR* zebrafish (1 year-old adults), with ablated region outlined in red dotted lines. Below shows in situ hybridization for *col1a1a (*magenta) and *col2a1a (*green) in adjacent sections, with *col2a1a* marking new chondrocytes and *col1a1a* upregulated in the adjacent periosteum. **e,** Diagram of the adult M before and after ablation showing new cartilage formation from the perichondrium. Scale bars = 20 um. aCh, anterior ceratohyal growth plate; pCh, posterior ceratohyal growth plate.

### New chondrocytes arise from the perichondrium following cartilage ablation

The finding that new cartilage arose around the entire ablated M, which as a permanent cartilage is surrounded by perichondrium, but only in perichondrium regions of the growth plates of ablated Ch and Pq endochondral bones, suggested that perichondrium may be the source of regenerative chondrogenesis. Further, in situ hybridization for the osteoblast marker *col1a1a* in ablated adult M showed upregulation in the periosteum but no morphological evidence of new chondrocytes (Fig. 2d,e). To confirm the perichondrium origin of regenerative chondrogenesis, we used a *hyal4* enhancer, identified as selectively accessible in perichondrium from single-cell RNA and accessible chromatin sequencing analysis of cranial neural crest-derived cells (Fabian et al., 2022), to generate a *hyal4:GFP* transgenic line. Whole-mount live imaging at 14 dpf and staining of sections with anti-GFP and anti-mCherry antibodies at 27 dpf confirmed restricted expression of *hyal4:GFP* in perichondrium surrounding M and Pq cartilages but not in *col2a1a^BAC^:mCherry-NTR*+ chondrocytes (Fig. 3a,b). Next, we treated *hyal4:GFP*; *col2a1a^BAC^:mCherry-NTR* fish with MTZ at 21 dpf and examined GFP and mCherry expression at 6 dpa in sections. In new mCherry+ chondrocytes around ablated M, we observed strong *hyal4:GFP* expression (Fig. 3c). To determine whether this was due to retention of GFP protein in new chondrocytes from their origin in the perichondrium, versus new induction of *hyal4:GFP*, we examined endogenous *hyal4* expression during cartilage regeneration. In DMSO-treated control animals, we observed mutually exclusive expression of *hyal4* in the perichondrium and *sox9a* in chondrocytes of M cartilage. At 6 dpa, *hyal4* was strongly expressed in the perichondrium yet only a few *sox9a*+ new chondrocytes displayed weak *hyal4* expression (Fig. 3d). Together, these data support *hyal4*+ perichondrium being the source of new chondrocytes following cartilage ablation.

**Figure 3:**
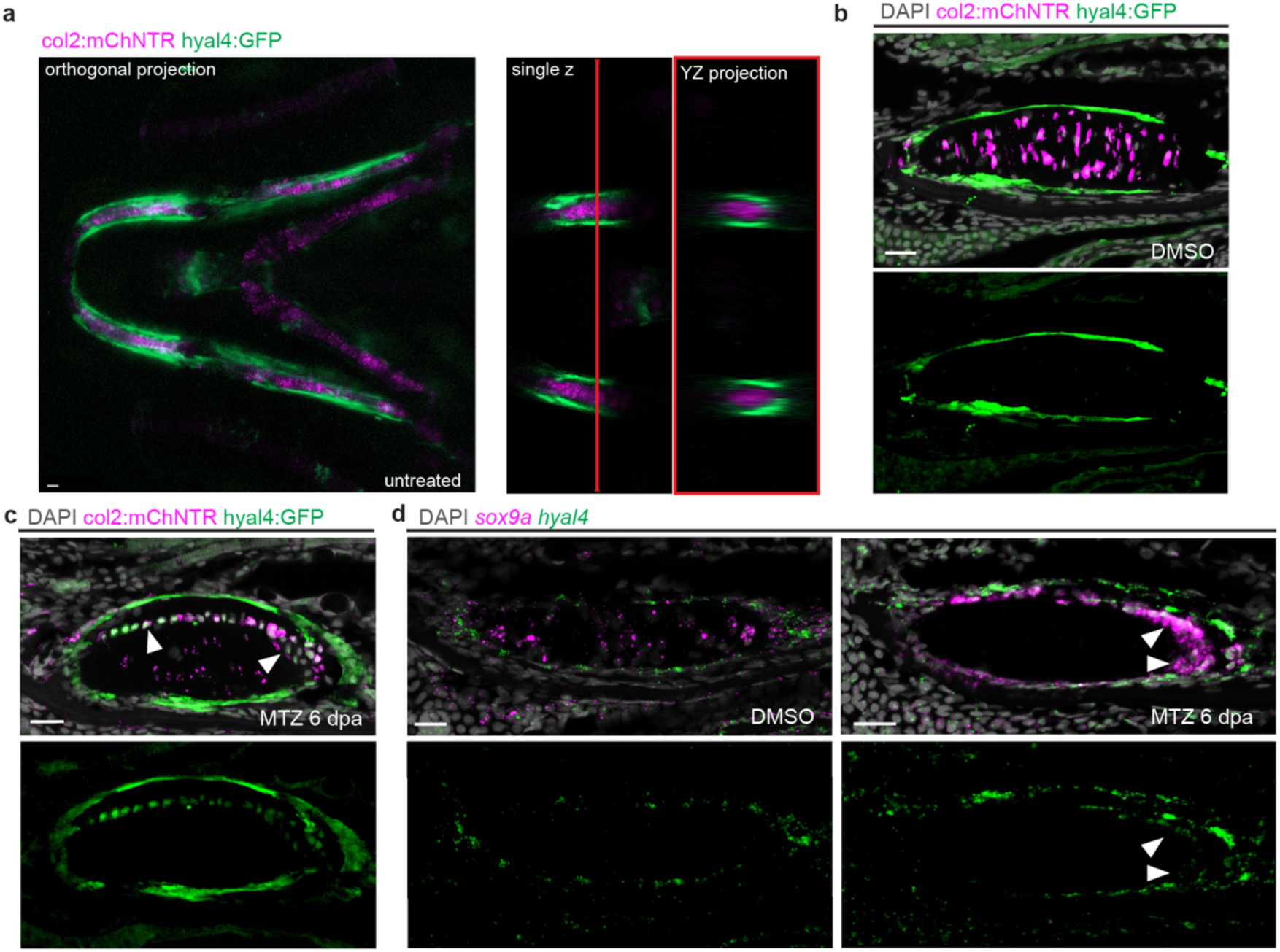
Perichondral origin of regenerative chondrocytes. **a,** Confocal imaging from a ventral view of the jaws of 14 dpf *hyal4:GFP*; *col2a1aBAC*:*mCherry-NTR* fish shows non-overlapping expression of *hyal4:GFP* within the perichondrium and *col2a1aBAC*:*mCherry-NTR* within the chondrocytes of M and Pq cartilages. To the right, single z-plane and YZ projection confirm absence of *hyal4:GFP* expression in chondrocytes. **b, c,** Sections of the M cartilages of 27 dpf *hyal4:GFP*; *col2a1aBAC*:*mCherry-NTR* fish treated with DMSO or MTZ and stained with anti-mCherry (magenta) and anti-GFP (green) antibodies. In DMSO controls, *hyal4:GFP* (green) is restricted to the perichondrium and not expressed in mCherry+ chondrocytes (magenta). In MTZ-treated animals at 6 dpa, new chondrocytes are double-positive for *hyal4:GFP* and *col2a1aBAC*:*mCherry-NTR* (white arrowheads). **d,** RNA expression of *sox9a* (magenta) and *hyal4* (green) in sections of M cartilages from 27 dpf fish. In DMSO-treated controls, *hyal4* expression in the perichondrium and *sox9a* expression in chondrocytes are mutually exclusive. In MTZ-treated fish at 6 dpa, *sox9a* is upregulated in newly forming chondrocytes which are largely negative for *hyal4* (arrowheads). Scale bars = 20 um.

### Dynamic retinoic acid pathway gene expression during cartilage regeneration

Given the morphological similarity of ablation-induced cartilage outgrowths to the exostoses seen in MO, and the putative role of RA signaling in this disease, we examined the expression of RA pathway genes following zebrafish cartilage ablation. Zebrafish have two RAR*α* and two RARγ receptor genes (*raraa*, *rarab*, *rarga*, *rargb*) but no RARβ orthologs. In DMSO-treated controls at 6 wpf, we observed weak expression of all four RARs in the perichondrium and cartilage of the Ch endochondral bone (Fig. 4a). At 3 dpa, all four RARs continued to be expression in the perichondrium, with only *rarga* and *rargb* upregulated in newly forming chondrocytes of Ch cartilage outgrowths (Fig. 4b). Aldehyde dehydrogenase family, member 2 (Aldh1a2) converts retinaldehyde to all-trans-retinoic acid (ATRA), which is then inactivated through hydroxylation by Cytochrome P450 26B1 (Cyp26b1). In DMSO-treated controls at 6 wpf, little expression of *aldh1a2* and *cyp26b1* was seen in M cartilage and the Ch endochondral bone (Fig. 4c,d). In contrast, ablation of chondrocytes induced strong upregulation of *aldh1a2* and *cyp26b1* at 3 and 4 dpa, which waned by 6 dpa (Fig. 4c,d). Whereas *aldh1a2* was largely restricted to the perichondrium at these stages, *cyp261* expression was enriched in *col2a1a*+ chondrocytes within the cartilage outgrowths at 4 dpa. While the expression of the RA synthesis gene *aldh1a2* and all four RARs in the perichondrium of outgrowths may suggest roles of RA-dependent signaling in the initiation of chondrogenesis, the expression of the RA degradation gene *cyp26b1* and *rarga/rargb* in newly forming chondrocytes may suggest a shift to ligand-less repression by RARγ for chondrocyte maturation.

**Figure 4:**
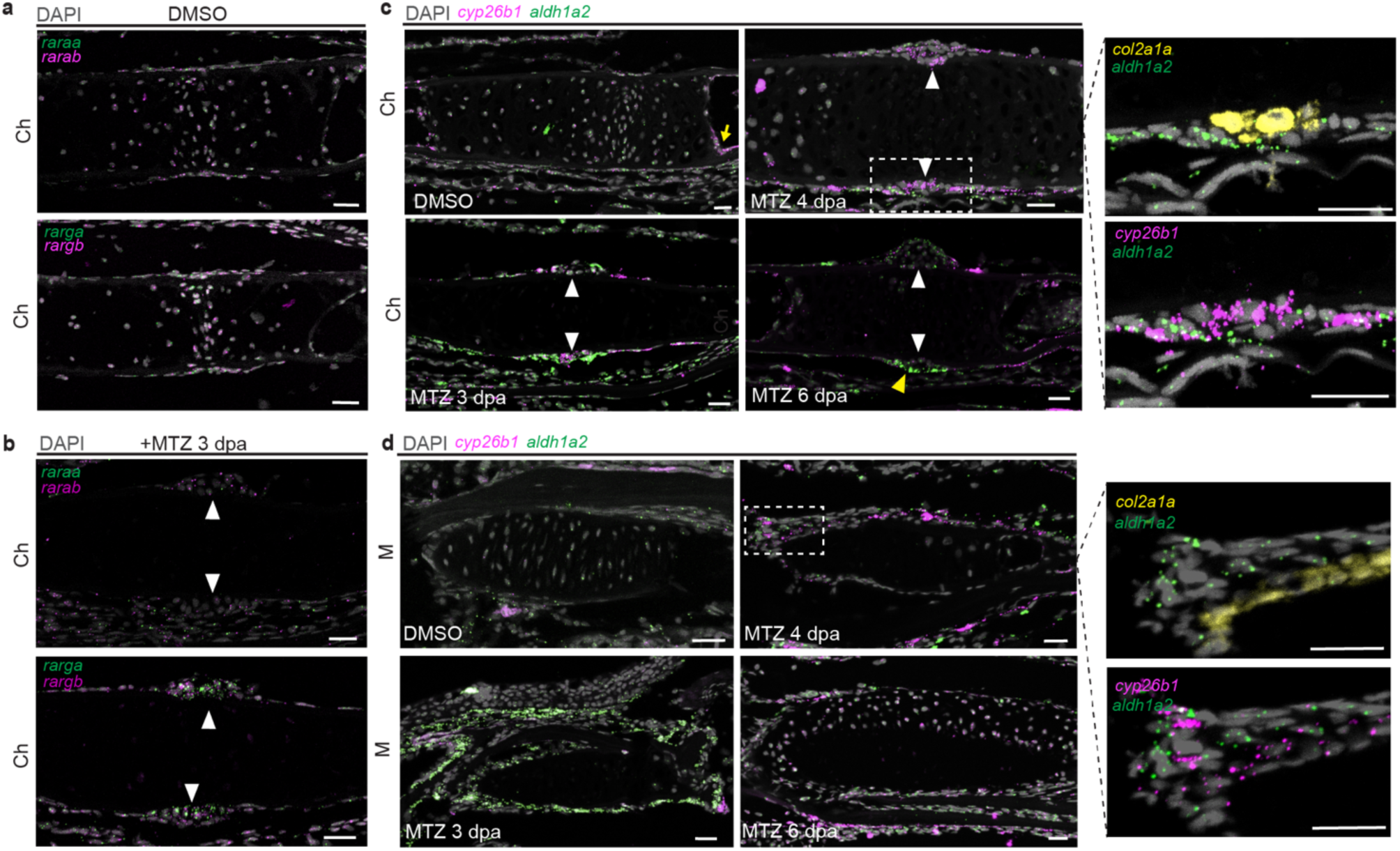
Retinoic acid pathway gene expression during cartilage regeneration. **a, b,** In situ hybridization for *raraa, rarab, rarga,* and *rargb* in sections of the Ch cartilages in juvenile (6-8 wpf) *col2a1aBAC*:*mCherry-NTR* fish treated with DMSO or MTZ and fixed at 3 dpa. DAPI labels nuclei in white and arrowheads denote cartilage exostoses in ablated fish. **c,d,** In situ hybridization for *cyp26b1* and *aldh1a2* in sections of the Ch and M cartilages in juvenile (6-8 wpf) *col2a1aBAC*:*mCherry-NTR* fish treated with DMSO or MTZ and fixed at 3, 4 or 6 dpa. DAPI labels nuclei in white and white arrowheads denote cartilage exostoses. Yellow arrow denotes *cyp26b1* expression in terminal hypertrophic zone of the Ch cartilage in DMSO controls, and yellow arrowhead denotes *aldh1a2* expression in perichondrium surrounding cartilage exostosis at 6 dpa. Dashed boxed regions are magnified to right and show segregation of *aldh1a2* to the perichondrium layer of exostoses and co-enrichment of *cyp26b1* and *col2a1a* in chondrocytes. Scale bars = 20um.

### Precise regulation of RA signaling is required for ablation-induced chondrogenesis

As expression data support ligand-less repression, in particular through RARγ, in new chondrocyte differentiation following ablation, we examined the effect of increasing RARγ signaling on cartilage regeneration. To do so, we treated *col2a1a^BAC^:mCherry-NTR*; *col2a1a1^BAC^:GFP* fish at 6-8 wpf with MTZ for 24 h to ablate chondrocytes and then exposed fish continuously to PVO, ATRA or DMSO control via daily water changes. Compared to 9/12 DMSO-treated control fish that displayed new *col2a1a1^BAC^:GFP*+ chondrocytes around M and the Ch growth plate, 3/3 PVO-treated and 3/3 ATRA-treated animals displayed a complete absence of new chondrocytes at 6 dpa that we confirmed by Alcian blue staining (Fig. 5a-c). TUNEL staining of apoptotic cells at 3 dpa ruled out that this inhibition was due to a prevention of MTZ-induced chondrocyte death (Supp. Fig. 1). In addition, loss of ablation-induced chondrogenesis in PVO-treated animals was associated with an upregulation of the osteoblast marker *col10a1a* in the perichondrium at 3 dpa (Fig. 5d). Compared to DMSO-treated animals where *col10a1a* marked the bone collar distant to Ch growth plates, *col10a1a* expression in MTZ-treated ablated animals expanded within the periosteum to abut the nascent exostoses but was excluded from the perichondrium. Upon PVO treatment of ablated animals, *col10a1a* formed a continuous expression domain throughout the periosteum and perichondrium that correlated with a lack of new cartilage formation. Lastly, given expression of *aldh1a2* and all four RARs in the perichondrium, we investigated the effect of inhibiting RA signaling on ablation-induced chondrogenesis. Consistent with RA signaling also be required for new chondrogenesis, daily treatment of ablated fish with the RA synthesis inhibitor DEAB or the RAR antagonist AGN193109 completely eliminated the appearance of new *col2a1a1^BAC^:GFP*+ chondrocytes around ablated M at 6 dpa (Fig. 5e-f). Our data therefore support both activating and inhibitory roles of RA signaling that are likely spatially segregated between the perichondrium and differentiating chondrocytes of ablation-induce cartilage outgrowths.

**Figure 5:**
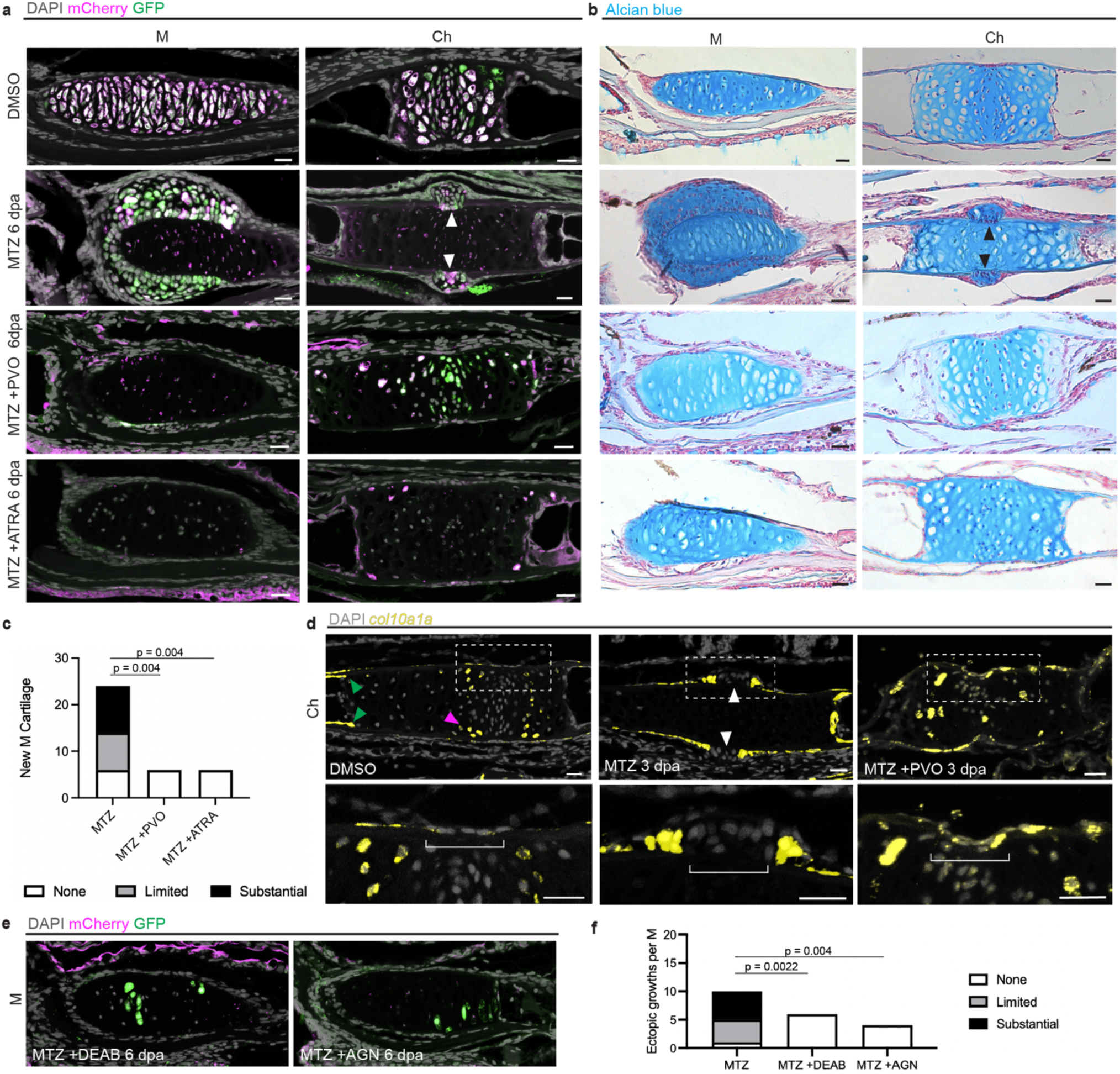
Precise RA signaling levels are required for cartilage regeneration. **a,b,** Juvenile (6-8 wpf) *col2a1aBAC*:*mCherry-NTR; col2a1aBAC*:*GFP* fish were treated with DMSO or MTZ for 24 hours, with MTZ-treated fish then treated for 6 days with DMSO control, 0.2 uM PVO, or 0.1 uM ATRA. At 6 dpa, new cartilage was assessed in sections of M or Ch by antibody staining with anti-mCherry (magenta) and anti-GFP (green) (**a**) or Alcian blue staining (**b**). Nuclei were visualized with DAPI in **a** and nuclear fast red in **b**. Arrowheads depict exostoses in fish treated with MTZ alone but not combined MTZ + PVO and MTZ + ATRA. **c,** Fraction of animals treated with MTZ alone or MTZ with PVO or ATRA showing no new M cartilage versus limited or substantial new cartilage. *p* values represent Fisher’s exact test. **d,** In situ hybridization for *col10a1a* in sections of the Ch cartilages in juvenile (6-8 wpf) *col2a1aBAC*:*mCherry-NTR* fish treated with DMSO or MTZ for 1 day and then incubated with DMSO or PVO for 3 days. DAPI labels nuclei in white. Green arrowheads denote *col10a1a* expression in osteoblasts of the bone collar, magenta arrowhead expression in hypertrophic chondrocytes, and white arrowheads denote cartilage exostoses. White dashed boxes are magnified below, with brackets indicating the perichondrium/exostoses regions in which *col10a1a* expression is expanded by PVO treatment. **e,** Juvenile (6-8 wpf) *col2a1aBAC*:*mCherry-NTR; col2a1aBAC*:*GFP* fish were treated with MTZ for 1 day, followed by 6 days with 10 mM DEAB or 0.2 uM AGN193109. At 6 dpa, antibody staining with anti-mCherry (magenta) and anti-GFP (green) on sections of M cartilages revealed an absence of exostoses. Note a few GFP+ chondrocytes escaped ablation within M cartilages. Scale bars = 20 um. **f,** Fraction of animals treated with MTZ alone or MTZ with DEAB or AGN193109 showing no new M cartilage versus limited or substantial new cartilage. *p* values represent Fisher’s exact test.

### Requirement of BMP signaling in cartilage regeneration

BMP signaling has known roles in cartilage formation during development, and stimulation of RARγ signaling with PVO has been suggested to inhibit BMP signaling in the context of MO (Inubushi et al., 2017). We therefore investigated the role of BMP signaling in ablation-induced chondrogenesis and its interaction with RA signaling. In DMSO-treated *col2a1a^BAC^:mCherry-NTR* fish at 6 wpf, we observed expression of the BMP target gene *id1* in growth plate chondrocytes and perichondrium of Ch (Fig. 6a). Three days after MTZ-induced cartilage ablation, *id1* expression was upregulated in newly forming chondrocytes adjacent to Ch growth plates (Fig. 6a). Next, we sought to determine if BMP inhibition would prevent ablation-induced regeneration. To compare the effects of BMP inhibition with RARγ activation, we treated juvenile (6-8 wpf) *col2a1a^BAC^:mCherry-NTR* fish with MTZ for 24 h to ablate cartilage and then subjected fish to daily water changes containing the BMP receptor antagonist LDN189193, PVO, or DMSO control (Fig. 6b). At 6 dpa, we observed substantial new cartilage around 6/6 ablated M cartilages in DMSO-treated controls, and no new cartilage in 3/4 PVO-treated M cartilages. Treatment with LDN189193 resulted in a milder inhibition of new chondrogenesis, with 2/7 M cartilages displaying substantial new cartilage, 3/7 displaying limited new cartilage, and 2/7 failing to mount a chondrogenic response (Fig. 6b,c). We also used TUNEL cell death staining to confirm that LDN189193 treatment did not prevent MTZ-induced chondrocyte ablation (Supp. Fig. 1). To further compare the mechanisms by which LDN189193 and PVO inhibit ablation-induced chondrogenesis, we performed in situ hybridization for *sox9a* and *id1* at 3 dpa (Fig. 6d). Compared to DMSO-treated animals in which *sox9a* and *id1* were strongly co-expressed in the perichondrium of ablated M cartilage, LDN189193-treated animals had severe reduction of *id1* expression, consistent with BMP inhibition. In contrast, PVO-treated animals had milder reduction of *id1* expression. In addition, PVO-treated but not LDN189193-treated animals had a complete loss of *sox9a* expression in M perichondrium, suggesting that RA activation suppresses the initiation of chondrogenesis while BMP inhibition may act later to inhibit the subsequent differentiation and/or expansion of chondrocytes.

**Figure 6:**
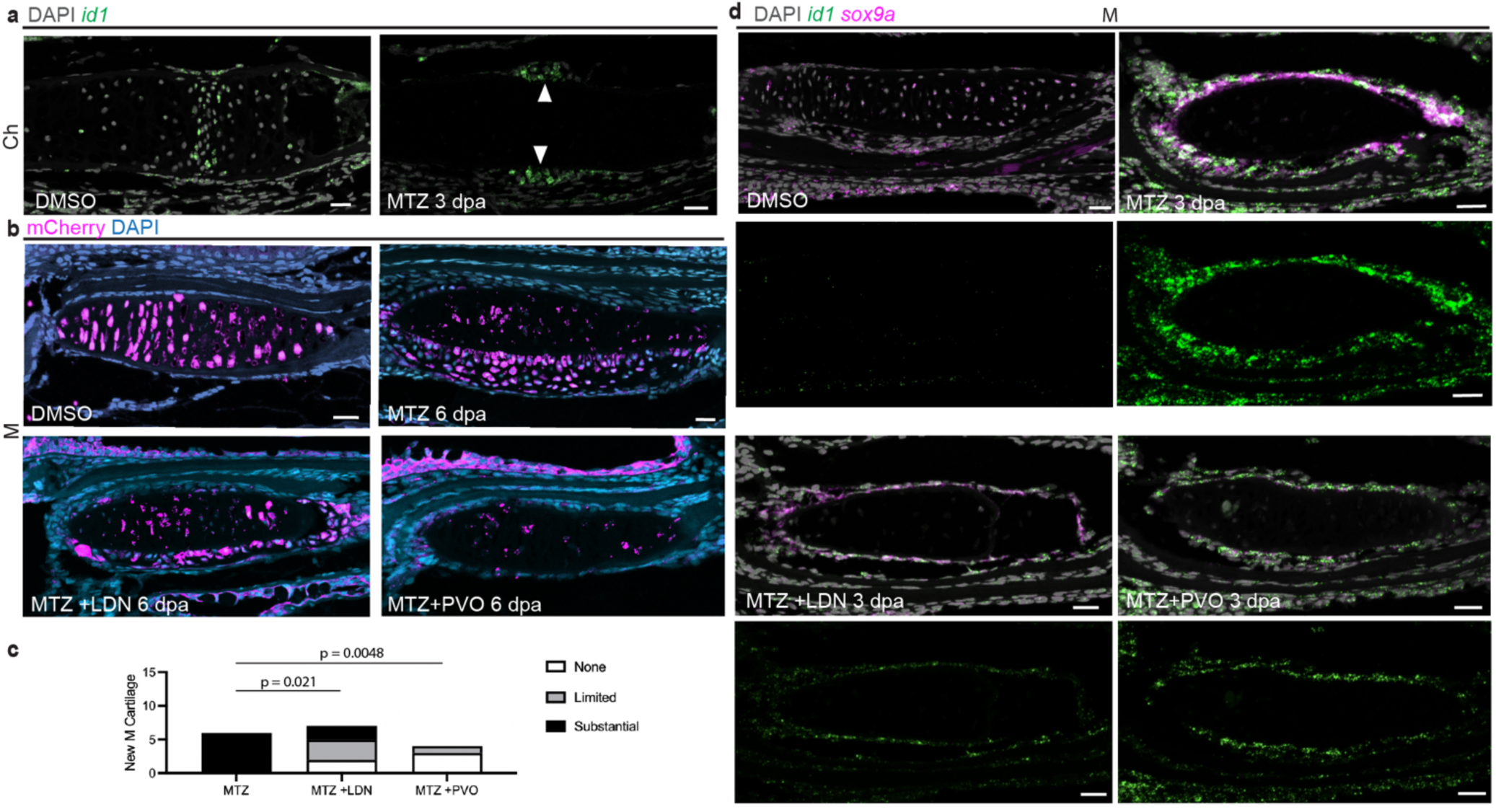
BMP inhibition impacts cartilage regeneration. **a,** In situ hybridization for *id1* in sections of the Ch cartilages of juvenile (6-8 wpf) *col2a1aBAC*:*mCherry-NTR* fish treated with DMSO or MTZ for 1 day and then fixed for analysis 3 days later. DAPI labels nuclei in white, and arrowheads indicate exostoses. **b,** Juvenile (6-8 wpf) *col2a1aBAC*:*mCherry-NTR; col2a1aBAC*:*GFP* fish were treated with MTZ for 1 day, then treated for 3 days with DMSO only, 1 uM LDN-193189, or 0.4 uM PVO, and then fixed for analysis after an additional 3 days. Anti-mCherry antibody staining (magenta) on sections of M cartilages revealed new cartilage in MTZ-treated animals that was blocked by LDN-193189 or PVO. **c,** Fraction of animals treated with MTZ alone or MTZ with LDN-193189 or PVO showing no new M cartilage versus limited or substantial new cartilage. *p* values represent Fisher’s exact test. **d,** In situ hybridization for *id1* and *sox9a* in sections of the M cartilages of juvenile (6-8 wpf) *col2a1aBAC*:*mCherry-NTR* fish treated with DMSO or MTZ for 1 day, and then DMSO only, 1 uM LDN-193189, or 0.4 uM PVO for 3 days. Merged images are shown above with *id1* channel alone shown below. Scale bars = 20um.

### RA signaling promotes chondrocyte exit from the growth plate

A major side effect of PVO in mice and adolescent patients is accelerated closure of the growth plates (Inubushi et al., 2018; Pignolo et al., 2023). We find that this effect of PVO on growth plates is conserved in zebrafish. When juvenile (6-8 wpf) zebrafish are treated continuously for 6 days with PVO and then fixed, sectioned, and stained with Alcian blue to detect cartilage, we observed a severe loss of growth plate cartilage in the endochondral Ch bone (Fig. 7a,b). This effect was most pronounced in the anterior Ch where 4/5 PVO-treated growth plates had no detectable cartilage remaining after treatment. Similar though milder effects were seen upon treatment for 6 days with ATRA, but not with the RA synthesis inhibitor DEAB (Fig. 7a,b). To understand how PVO affects growth plate architecture at earlier stages, we performed RNAscope in situ hybridization for markers of growth plate zones in juveniles treated for 3 days with PVO. Whereas we observed severe reductions in the number of cells expressing pre-hypertrophic and hypertrophic markers *ihha* and *col10a1a* in PVO-treated animals, the number of cells expressing the terminal hypertrophic marker *mmp13b* was increased (Fig. 7c). Based on background fluorescence from the DAPI channel, which we used to label nuclei, we observed a degradation of the cartilage matrix at the edges of the growth plate in PVO-treated animals, giving the appearance of many chondrocytes breaking away from the growth plate. Further, we found that many of these cells expressed the pre-osteoblast (and hypertrophic) marker *runx2b* (Fig. 7c). To confirm that the cells accumulating in the marrow space of PVO-treated animals were derived from growth plate chondrocytes, we treated *col2a1a-R2:CreERT2*; *bactin:B>R* fish with 4-OH-tamoxifen at 5 dpf to permanently label chondrocytes, raised these fish to juvenile stages (6 wpf), and then incubated with PVO for 3 days. Compared to DMSO-treated controls, we observed an increase in loosely packed chondrocyte lineage cells at the edges of PVO-treated Ch growth plates (Fig. 7d,e). These results suggest that stimulating RA signaling, in particular through RARγ, results in the accelerated terminal differentiation of hypertrophic chondrocytes and their exit from the growth plate into the marrow cavity (Fig. 7f).

**Figure 7:**
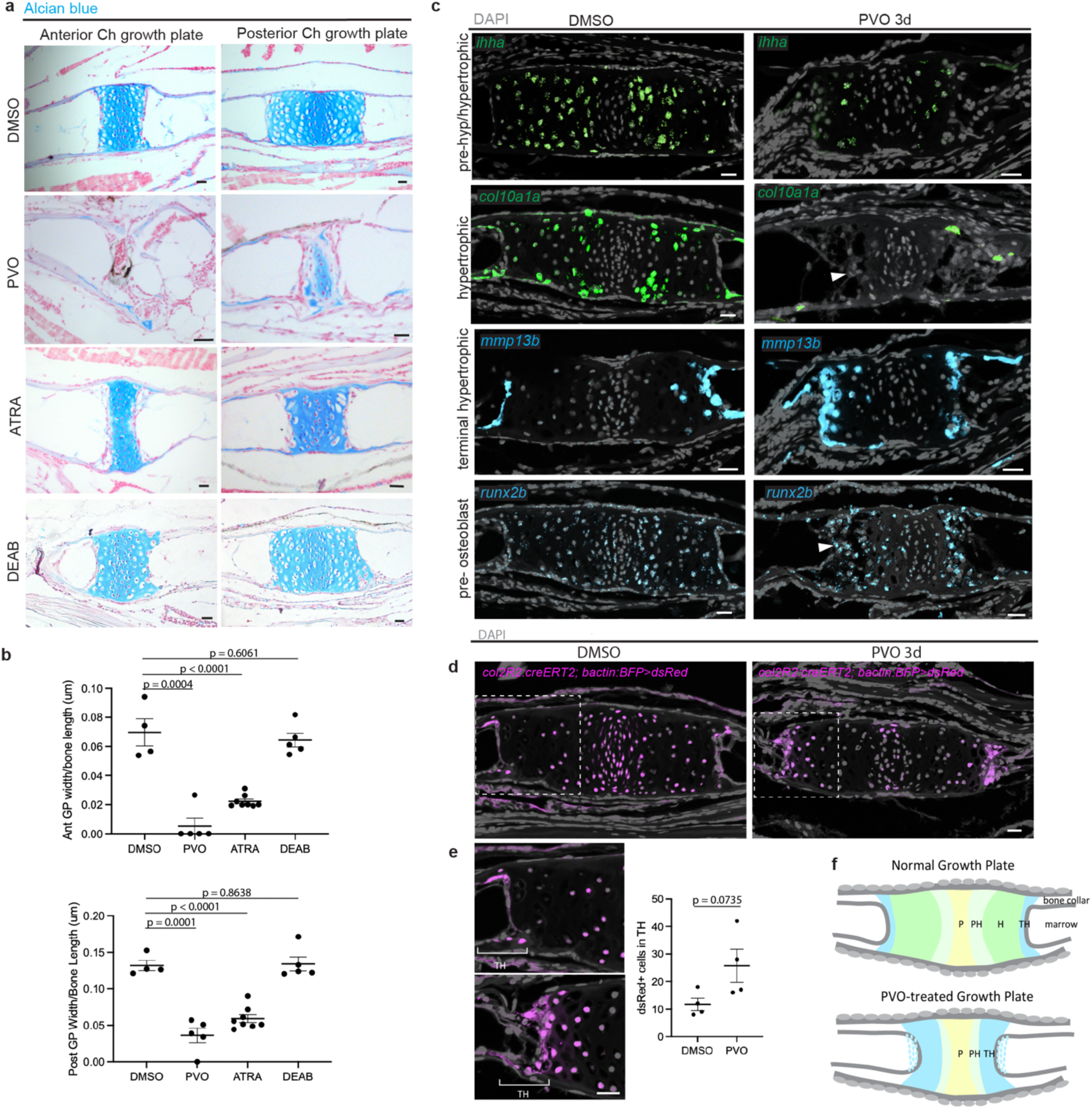
Retinoic acid activation induces growth plate collapse. **a,** Alcian blue-stained sections of the posterior and anterior Ch growth plates in juvenile (6-8 wpf) fish continuously treated with DMSO, 0.2 uM PVO, 0.1 uM ATRA or 10 mM DEAB for 6 days. Nuclei are labeled with nuclear fast red. **b,** The width of the anterior and posterior growth plate (GP) relative to total Ch bone length are shown for each condition in **a**. Standard errors of the mean are shown, with *p* values calculated with a student’s t-test. **c,** In situ hybridization in sections of the posterior growth plate of the Ch cartilages of juvenile (6-8 wpf) fish treated with DMSO or 2 uM PVO for 3 days. Probes include markers of the pre-hypertrophic and hypertrophic zones (*ihha*), hypertrophic zone (*col10a1a*), terminal hypertrophic zone (*mmp13b*), and pre-osteoblasts (*runx2b*). Arrowheads denote apparent breakdown of the cartilage matrix, visualized by weak DAPI signal that also labels nuclei more strongly. **d,e,** Lineage tracing of chondrocytes was performed by overnight treatment of *col2a1a-R2-E1b:CreERT2*; *bactin2:loxP-BFP-loxP-dsRed* fish with 4-OHT at 5 dpf. The same fish were then treated at juvenile stages (6-8 wpf) with DMSO or 0.2 uM PVO for 3 days. Anti-dsRed antibody staining on sections of the posterior growth plates of Ch cartilages reveals chondrocyte lineage cells in magenta. Boxed regions are magnified in **e** to show increased numbers of chondrocyte lineage cells in the terminal hypertrophic zone and marrow cavity in PVO-treated fish versus DMSO-treated controls, which is quantified to the right. Standard errors of the mean are shown, and the *p* value was calculated with a student’s t-test. Scale bars = 20um. **f,** Model of how activation of retinoic acid signaling by PVO alters growth plate architecture, including increased numbers of chondrocyte lineage cells (blue dots) entering the marrow cavity. P, proliferative zone; PH, prehypertrophic zone; H, hypertrophic zone; TH, terminal hypertrophic zone.

## Discussion

Here we show that zebrafish are able to broadly regenerate cartilage throughout their lifetime, albeit imperfectly with new cartilage forming around remnant dead cartilage. Imperfect regeneration has also been observed in lizard tails, where the bony vertebrae are replaced by an unsegmented cartilage tube (Lozito and Tuan, 2015). Further, ablation-induced cartilage regeneration in zebrafish shares many features with MO, including a likely perichondrium origin and dependence on precisely regulated RA and BMP signaling. We also uncover a conserved role for RA signaling in regulation of growth plate maturation in zebrafish and mouse, with RA signaling promoting the terminal differentiation and exit of hypertrophic chondrocytes into the marrow cavity. The ability of RA signaling to suppress the chondrocyte state in both growth plate and regenerating cartilage may thus explain the growth plate closure defects seen when treating MO and other diseases of heterotopic ossification with RA agonists.

### Similarities of ablation-induced cartilage regeneration and diseases of heterotopic ossification

In addition to MO, heterotopic ossification (HO) resulting from trauma (DEY et al., 2017) and Fibrodysplasia Ossificans Progressiva (FOP), an extremely debilitating disease of soft tissue ossification, proceed through a cartilage intermediate. Although the role of injury in MO is less clear, it is possible that the cartilage regeneration induced by ablation in zebrafish involves a similar type of injury signal to that in HO and FOP, such as recruitment of immune cells to the ablated cartilage. Alternatively, chondrocyte ablation in zebrafish may result in alterations to the proteoglycan matrix that mimic the reduction of HSPGs caused by *EXT1* or *EXT2* heterozygosity in MO patients. Previous studies using C-14 radiolabeling of human articular cartilage revealed that while collagen proteins are very long-lived, proteoglycans and their associated glycosoaminoglycans turn over rapidly in days (Heinemeier et al., 2016). In culture, treatment with a small molecule HSPG antagonist increases chondrocyte differentiation which can be blocked by treatment with PVO (Pacifici, 2018). One possibility is that cells in the perichondrium sense chondrocyte health by the presence of sufficient HSPGs. When chondrocytes are ablated and can no longer produce new HSPGs, or when HSPG synthesis genes such as *EXT1* and *EXT2* are mutated, perichondrium cells would sense this as a cartilage deficiency and initiate chondrogenesis as a repair response. This could be seen as hijacking of normal mechanisms to restore injuries to growth plate cartilage during fetal and adolescent development. MO could therefore be considered a disease of misregulated cartilage regeneration. The extent to which cartilage ablation affects HSPG levels in zebrafish will need to be formally tested, as our attempts to use antibodies to monitor HSPGs following ablation were unsuccessful.

### Distinct roles of RA and BMP signaling in ablation-induced cartilage regeneration

Following cartilage ablation, we observed upregulation of the *rarga* and *rargb* receptor genes, the RA synthesis gene *aldh1a2*, and the RA degradation gene *cyp26b1* in the ectopic cartilage outgrowths. Enrichment of *aldh1a2* in the perichondrium surrounding the outgrowths and *cyp26b1* in the newly forming chondrocytes suggests that RA levels may be higher in the perichondrium than in chondrocytes. As *cyp26b1* can be an RA target gene, it may be that RA signaling promotes chondrocyte initiation in the perichondrium, followed by *cyp26b1* upregulation and RA degradation to promote ligand-less RARγ repression in newly forming chondrocytes. Consistent with dynamic RA signaling being required for ablation-induced chondrogenesis, both activation and inhibition of RA signaling blocked cartilage regeneration. Unfortunately, we were unable to directly monitor RA activity during cartilage regeneration using the previously published RA reporter lines *Tg(12XRARE-ef1a:gfp)^sk72^* and *Tg(β-actin:GDBD-RLBD)^cch1^* as we found they became largely silenced by juvenile stages (D’Aniello et al., 2013; Waxman and Yelon, 2011). One role of ligand-less RARγ repression may be inhibition of periosteum/osteoblast genes such as *col10a1a*, as activation of ligand-dependent RARγ signaling by PVO resulted in expanded *col10a1a* expression throughout the domain that would normally form exostoses. Consistently, elevation of RA signaling in *cyp26b1* mutant zebrafish leads to premature and excessive ossification of endochondral bones (Laue et al., 2008; Spoorendonk et al., 2008). Thus, RARγ signaling may shift the perichondrium to an osteogenic periosteum state following ablation, preventing cartilage regeneration.

In addition to RA signaling, BMP signaling has been a major focus of study in heterotopic ossification, in particular due to the discovery that activating mutations in ACVR1/BMPR1 cause FOP. BMP signaling also has important roles in endochondral ossification, acting as an instructive signal at multiple stages of chondrogenesis, including during initial condensation, proliferative expansion, and hypertrophy (De Luca et al., 2001; Yoon et al., 2006). Sox9 was previously thought to be downstream of BMP, due to its upregulation after BMP bead implantation in chick limbs (Healy et al., 1999). However, in developing murine digits, *Sox9* is upregulated prior to *Bmpr1b* expression and its upregulation cannot be inhibited by exogenous addition of *Noggin* (Chimal-Monroy et al., 2003). In the *Fsp1-Cre*; *Ext1^fl/fl^* mouse model of MO, BMP signaling is upregulated in the perichondrium based on phospho-Smad1/5/8 staining (Inubushi et al., 2018, 2017), and PVO treatment decreases phospho-Smad1/5/8 levels (Inubushi et al., 2018; Pacifici, 2018). During ablation-induced cartilage regeneration in zebrafish, we also detect BMP activity in the perichondrium based on *id1* expression, and treatment with the BMP inhibitor LDN189193 partially prevented cartilage outgrowths. Whereas treatment of zebrafish with PVO following cartilage ablation also modestly decreased expression of *id1*, we observed that PVO but not LDN189193 resulted in severely reduced expression of *sox9a*, a master regulator of chondrocyte differentiation. Our data therefore support distinct roles of RA and BMP signaling in the initiation and expansion of ectopic cartilage growths, respectively.

It remains unclear how RA and BMP signaling are initiated in response to cartilage ablation. Although RA has not been documented to interact with HSPGs, some studies have suggested that HSPGs may sequester BMP ligands which are then released in response to ablation or injury (Huegel et al., 2013; Ruppert et al., 1996). It is also unknown why the periosteum does not initiate a chondrogenic response to cartilage ablation, despite cells in this layer being in close proximity to ablated hypertrophic chondrocytes. In response to bone fractures, the periosteum generates a cartilage callus that remodels to bone. However, the cartilage formed in response to bone fracture differs from that formed in response to cartilage ablation by co-expressing a hybrid chondrocyte-osteoblast gene signature (Kuwahara et al., 2019; Paul et al., 2016). These findings indicate that the perichondrium senses cartilage injuries and the periosteum senses bone injuries distinctly, mounting a pure cartilage versus an endochondral response, respectively. Understanding the molecular bases for the differential competencies of the perichondrium and periosteum will be a fruitful avenue of future investigation.

### RA signaling promotes the terminal differentiation and growth plate exit of hypertrophic chondrocytes

Treatment of heterotopic ossification with PVO has been associated with growth plate closure in adolescents, indicating roles of RA signaling in growth plate biology. For example, a phase 3 clinical trial of PVO to treat FOP revealed premature growth plate closure in 39.6% of adolescents (Pignolo et al., 2023). Our analysis of PVO effects on zebrafish growth plates reveals potential mechanisms of RA action. In PVO-treated growth plates, we observed a loss of *col10a1a*+; *ihha*+ hypertrophic chondrocytes and a concomitant increase in *mmp13b*+ terminal hypertrophic chondrocytes. Lineage tracing of chondrocytes in PVO-treated fish revealed increased exit of hypertrophic chondrocytes from the growth plate into the marrow cavity, which was associated with accelerated cartilage matrix breakdown and eventual complete collapse of the growth plate. Cells exiting the growth plate also expressed the pre-osteoblast marker *runx2b*, suggesting that RA signaling may enhance the chondrocyte to osteoblast transition. These findings are consistent with previous cell culture studies in chjck. Treatment of cultured chicken growth plate chondrocytes with RA enhanced their mineralization and alkaline phosphatase activity (Descalzi Cancedda et al., 1992; Iwamoto et al., 1994; Koyama et al., 1999; Wu et al., 1997), and PVO was found to suppress expression of the hypertrophic chondrocyte marker *Col10a1* in cultured mouse growth plate chondrocytes (Pacifici, 2018). One model is that blood vessels in the forming marrow cavity bring RA or its precursors to the late hypertrophic zone, which induces their terminal differentiation, including loss of chondrocyte identity and exit from the growth plate. Consistently, we observe enriched expression of the RA target gene *cyp26b1* at the edges of the zebrafish growth plates. Upon treatment with PVO, chondrocytes would undergo RA-dependent signaling to induce terminal hypertrophic maturation throughout the growth plate, leading to rapid collapse. Our findings therefore link RA-mediated suppression of chondrocyte identity in the injured perichondrium and terminal hypertrophic zone of the growth plate, thus providing an explanation for why targeted activation of RA signaling can both inhibit heterotopic ossification and cause growth plate closure.

## Methods

### Zebrafish Lines

The Institutional Animal Care and Use Committee of the University of Southern California approved all animal experiments (Protocol 20771). Experiments were performed on zebrafish (*Danio rerio*) of mixed AB/Tubingen background. Lines include *Tg(col2a1aBAC:GFP)^el483^* and *Tg(col2a1aBAC:mCherry-NTR)^el559^* (Askary et al., 2015), *Tg(col2a1a-R2-E1b:CreERT2)^e712^*(Giovannone et al., 2019), and *Tg(bactin2:loxP-BFP-loxP-DsRed)^sd27^*(Kobayashi et al., 2014). To generate *Tg(hyal4:GFP)^el866^*, we synthesized using gBlocks (Integrated DNA Technologies) a perichondrium-specific snATAC-seq peak for *hyal4* (chr25:27624853-25:27625759, GRCz11) (Fabian et al., 2022) and cloned it into a modified pDest2AB2 construct containing an E1b minimal promoter, GFP, and an eye-CFP selectable marker using In-Fusion cloning (Takara Bio). We injected plasmid and Tol2 transposase RNA (30 ng/μL each) into one-cell stage zebrafish embryos, raised these animals, and screened for founders based on GFP expression in the progeny. Injected animals were raised and outcrossed to identify stable germline founders. Of note, the eye-CFP selectable marker segregates independently from the *Tg(hyal4:GFP)^e866^* transgene (likely reflecting multi-copy integration) and thus was not used to maintain this line.

### Histology

All samples were prepared by fixation in 4% paraformaldehyde overnight. Fixed animals were washed 2× in phosphate-buffered saline with Tween® 20 (PBST) for 30 min then decalcified for one to two weeks with rocking in 20% EDTA at room temperature if over 14 dpf. The tissue was dehydrated through an ethanol and Hemo-De series and then embedded in paraffin. Paraffin embedding was in Paraplast X-tra (VWR, 15159-486). We cut blocks into 5-6 μm sections on a Shandon Finesse Me+ microtome (cat. no. 77500102) with High-Profile Disposable Surgipath DB80 HS Blades (VWR, 10015-022), and collected sections on Apex Superior Adhesive slides (Leica Microsystems, cat. no. 3800080).

Pentachrome and Trichrome staining were performed according to manufacturer’s instructions (Movat-Russell modified pentachrome stain kit, Newcomer Supply cat. no. 9150A; Gomori One-Step, Aniline Blue, trichrome stain kit, Newcomer Supply cat. no. 9176A). Alcian blue (Alcian blue powder 8GX, Sigma, A3157) staining on sections was performed by deparaffinizing slides, submerging slides in Alcian blue solution, pH 1.0 (1g of powder in 100mL ddH2O, pH adjusted with HCl) for 15 min, thoroughly rinsing with DI water (5x 1 min rinses), followed with a counterstain of nuclear fast red (Vector, H-3403) for 25 seconds before rinsing thoroughly again with DI water (5 x 1 min rinses). The slides were dehydrated again through subsequent washes of 100% EtOH (1 x 4 min), Hemo-De (1 x 4 min) and finally xylene (1 x 4min) before mounting with Cytoseal (VWR, 48212-154).

### RNA In Situ Hybridization

RNAscope probes were synthesized by Advanced Cell Diagnostics in channels 1 through 4. Channel 1 probes: *col10a1a, ihha, id1, col2a1, rarga*. Channel 2 probes: *cyp26b1, sox9a, runx2b, raraa*. Channel 3 probes: *sox9a, aldh1a2, rarab*. Channel 4 probes: *hyal4, mmp13, rargb*. Paraformaldehyde-fixed paraffin-embedded sections were deparaffinized, and the RNAscope Fluorescent Multiplex v2 Assay was performed according to manufacturer’s protocols with the ACD HybEZ Hybridization oven. Specific modifications to the RNAscope protocol included using a 4 min steaming step (IHC World, IW-1102) during target retrieval and an extended 30 min protease step using the provided Protease Plus reagent. Slides were mounted with DAPI Fluoromount-G Mounting Solution (Southern Biotech, 0100-20). For fluorescent TSA in situ hybridization (*col1a1a and col2a1a* in Fig. 2d; *col10a1a* in Fig. 7c), paraffin sections were deparaffinized, permeabilized for 5 min with 7.5 ug/mL proteinase K solution, hybridized with probes at 68 °C overnight, and blocked with Perkin Elmer blocking powder (FP1020). Digoxigenin-labeled or dinitrophenol-labeled riboprobes were detected with anti-DIG-HRP (1:500, Perkin Elmer, NEF832001EA) or anti-DNP-HRP (1:200, Akoya, FP1129), respectively, and visualized using TSA Fluorescein (1:75, Akoya, NEL701A001KT), TSA Cy3 (1:75, Akoya, NEL704A001KT), or TSA Cy5 (1:100, Akoya, NEL705A001KT). The *col1a1* riboprobe was generated by PCR amplification of cDNA with primers 5′-GTATTGTAGGTCTCCCTGGACAAA-3′ and 5′-TGTTCTTGCAGTGGTATGTAATGTT-3′, the *col2a1a* riboprobe with primers 5′-GTAAAGATGGAGAGACTGGACCTTC-3′ and 5′-ATTCTCTCCTCTGTCTCCCTGTTT-3’, and the *col10a1* riboprobe with primers 5′-GTCTAAAAGGTGACAGAGGAGTACCT-3′, and 5′-GTAGACACTGATCAGTAACAAGGAAACA-3′. Amplified products were cloned into pCR-BluntII-TOPO. pCR-BluntII-TOPO plasmids were linearized by restriction digestion (enzyme dependent on direction of blunt insertion and probe sequence), and RNA probes were synthesized using either T7 polymerase (Roche, 10881775001) or Sp6 polymerase (Roche, 11487671001) depending on direction of blunt insertion.

### Immunohistochemistry and TUNEL Staining

To detect mCherry or GFP on sections, paraformaldehyde-fixed paraffin-embedded sections were deparaffinized followed by a 7 min −20 °C 100% acetone antigen retrieval step. For immunohistochemistry of BrdU on sections, antigen retrieval was performed by treating slides with citrate buffer (pH 6.0) in a steamer set (IHC World, IW-1102) for 20 min, subsequently removing slides from the steamer, and allowing to cool for an additional 20 min. Blocking was for 30 min at room temperature with PBST/1%DMSO/2% normal goat serum (Jackson ImmunoResearch, 005-000-121). Primary antibodies were applied overnight at 4 °C while secondary antibodies were applied for 1 hour at room temperature. Primary antibodies included rabbit anti-mCherry (1:100, Novus Biologicals, NBP2-25157), chicken anti-GFP (1:500, Abcam, ab13970), and mouse anti-BrdU (1:100, BioRad, MCA2060GA). Secondary antibodies included goat anti-rabbit Alexa Fluor 568 (1:500, Thermo Fisher, A11036), goat anti-rabbit Alexa Fluor 488 (1:500, Thermo Fisher, A11008), goat anti-chicken Alexa Fluor 488 (1:500, Thermo Fisher, A11039), and goat anti-chicken Alexa Fluor 568 (1:500, Thermo Fisher, A11041). Slides were mounted with DAPI Fluoromount-G Mounting Solution (Southern Biotech, 0100-20). To detect dying cells with TUNEL stain, slides were deparaffinized and rehydrated in 1x PBS, treated with proteinase K (7.5 ug/mL, 7 min), and followed by a postfix (4% PFA, 1x PBS; 20 min). Slides were then washed twice with 1x PBS for 5 min each. We then used the Fluorescein In situ Cell Death Detection Kit (Sigma, 11684795910), with reaction mixture applied for 60 min in the dark at 37 °C. Slides were then washed twice with 1x PBS for 5 min each before mounting.

### Drug Treatments

To permanently label chondrocytes with CreER-mediated recombination, *Tg(col2a1a-R2-E1b:CreERT2)^e712^*; *Tg(actab2:loxP-BFP-STOP-loxP-dsRed)^sd27^* fish were treated overnight at 5 dpf with 5 μM (Z)−4-Hydroxytamoxifen (4-OHT) (Sigma-Aldrich H7904) in EM. Frozen 4-OHT aliquots were heated for 10 min at 65°C prior to use to promote *trans* formation. Fish were then screened for dsRed+ conversion at 7 dpf using a Leica fluorescent stereomicroscope. For cartilage ablation, metronidazole (MTZ, Sigma-Aldrich, M1547) solutions (10mM MTZ, 0.5% DMSO in system water or EM) were made fresh and protected from light. *Tg(col2a1aBAC:mCherry-NTR)^el559^* animals were placed in MTZ solution for 24 hours in the dark before being rinsed with fresh system water or EM. Sacrifice was performed at 3, 4 or 6 days “post-ablation”, meaning post-removal of treatment. Palovarotene (PVO, Sigma-Aldrich, T94543) solutions were prepared from frozen stock aliquots of 12.05 mM in DMSO to working concentration of 0.2, 0.4 or 2 uM in fish system water or EM. LDN-193189 hydrochloride (Sigma-Aldrich, SML0559) solutions were prepared from frozen stock aliquots of 10 mM in DMSO to a working concentration of 1 uM in fish system water or EM. All-trans retinoic acid (Sigma-Aldrich, R2625) solutions were prepared from frozen stock aliquots of 1.0 mM in DMSO to working concentration of 0.1 uM in fish system water or EM. DEAB (Sigma-Aldrich, D86256) solutions were prepared from frozen stock aliquots of 500 mM in DMSO to working concentration of 10 mM in fish system water or EM. AGN194319 (Sigma-Aldrich, SML2665) solutions were prepared from frozen stock aliquots of 10 mM in DMSO to working concentration of 0.2 uM in fish system water or EM. All stock solutions were kept at -20°C. Animals were placed in drug solutions for 3 or 6 days, in the dark, with fresh drug changes every day.

### Imaging

Confocal images of section fluorescent sections and live transgenic fish were captured on a Zeiss LSM800 microscope using ZEN software, with representative sections or maximum intensity projections shown as specified. Brightness and contrast were adjusted in Adobe Photoshop with similar settings for experimental and control samples. Brightfield images of pentachrome, trichrome, and Alcian blue stains were acquired with a Leica DM 2500 microscope.

### Quantification and statistical analyses

We stage-matched control and experimental zebrafish by measuring standard body length (SL) from the tip of the snout to the edge of the hypuralia. Juvenile animals ranged from 15 mm - 20 mm. All transgenic patterns for (*Tg(col2a1aBAC:GFP)^el483^*, *Tg(col2a1aBAC:mCherry-NTR)^el559^*, and *Tg(hyal4:GFP)^el866^*) were seen in at least 8 independent animals. All in situ patterns were confirmed in at least 3 independent animals. Anti-mCherry and anti-GFP immunohistochemistry stains were performed in at least three animals, with the results used to assess amount of repair (contingency tables in Fig. 5-6). Statistical significance was determined by a student’s t-test for pair-wise comparisons (Fig. 7) or Fisher’s exact test for contingency tables (Fig. 5-6), using GraphPad’s Prism 10 software.

---

**Supplementary Figure 1:**
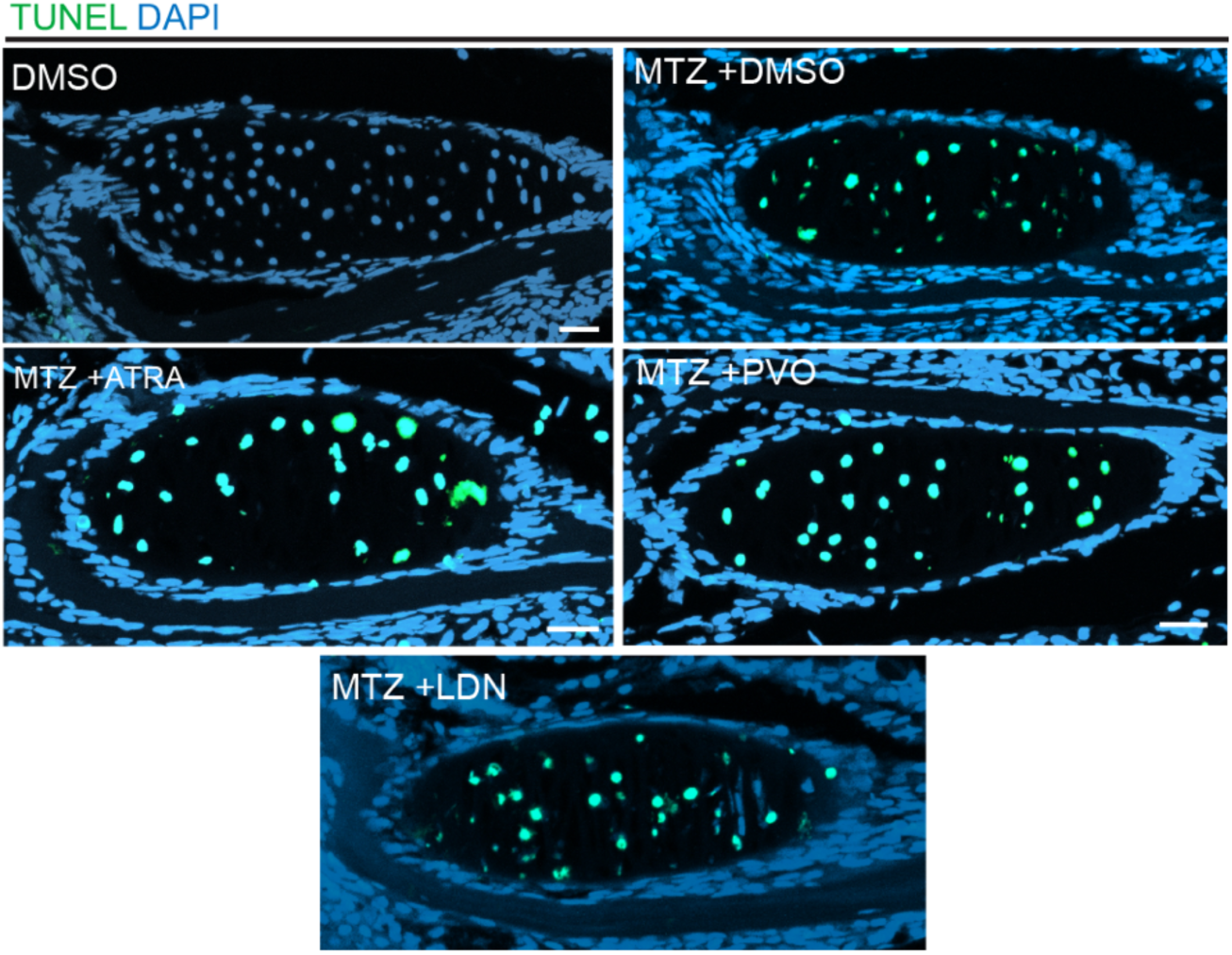
Drug treatments do not prevent MTZ-induced cartilage ablation. Sections through M cartilages of juvenile *col2a1aBAC*:*mCherry-NTR* fish (6-8 wpf) treated with DMSO (control) or 10mM MTZ for 24 hours to ablate cartilage. MTZ-treated animals were then treated for 3 consecutive days with daily water changes of either DMSO, 0.1 uM ATRA, 1 uM LDN-193189, or 0.4 uM PVO and fixed for processing. TUNEL staining labels apoptotic cells in green, and DAPI labels nuclei in blue. Scale bars = 20 um. MTZ, metronidazole; PVO, palovarotene; ATRA, all-trans retinoic acid; LDN, LDN-193189.

